# Evolutionary dynamics and molecular features of intra-tumor heterogeneity

**DOI:** 10.1101/214791

**Authors:** Franck Raynaud, Marco Mina, Giovanni Ciriello

## Abstract

The systematic assessment of intra-tumor heterogeneity is still limited and often unfeasible. *In silico* investigations of large tumor cohorts can be used to decipher how multiple clones emerge and organize into complex architectures. Here, we addressed this challenge by integrating mathematical modeling of cancer evolution with algorithmic inference of clonal phylogenies in 2,600 human tumors from 15 tumor types. Through numerical simulations, we could discriminate between observable and hidden intra-tumor heterogeneity, the latter characterized by clones that are missed by DNA sequencing of human samples. To overcome this limitation in human tumors, we show that population frequencies of detectable clones can be used to estimate the extent of hidden heterogeneity. Overall, simulated and human clonal architectures were highly concordant and showed that high numbers of clones invariably emerge through branching lineages. Interestingly, high numbers of alterations were not necessarily associated with high intra-tumor heterogeneity. Indeed, tumors with alterations in proliferation-associated genes exhibited high numbers of clonal mutations, but few clones. Instead, mutations of chromatin remodeling genes characterized tumors with high numbers of subclonal alterations and multiple clones. Our results identify evolutionary and genetic determinants of tumor clonal architectures to guide functional investigations of intra-tumor heterogeneity.

---

Cancer is a dynamic and ever-changing disease that mutates and evolves during its progression^1^. While the transformation from healthy to malignant cell is characterized by a few selected oncogenic alterations^2^, genetic instability in formed tumors promotes the acquisition of multiple lesions that diversify the cancer cell population^3^. As a result, each tumor is a composite of multiple *clones,* defined as groups of cells that are genetically identical within groups but different among them^4^.

Approaches based on single-cell profiles^5–8^ or multiple biopsies of the same tumor^9–11^ have revealed a daunting diversity among cancer cells. These observations questioned the ability of genomic studies of single tumor samples to provide a representative image of the disease. Unfortunately, single-cell analyses of tumors or profiling of multiple samples for each patient face technical and cost limitations, thus large scale datasets of these types are currently unavailable for systematic investigations. In response to these limitations, algorithmic approaches have been proposed to infer the clonal composition of a tumor from the genetic profile of a single sample^12–15^. Using such tools, different clonality and timing of emergence have been shown for specific therapeutically actionable mutations^16^ and an association between increasing intra-tumor heterogeneity and worse clinical outcome has been found^17^. However, these algorithmic approaches are limited in their ability to describe the dynamic emergence of intra-tumor heterogeneity and how this depends on features such as the rate of cell proliferation or acquisition of new alterations. From a theoretical perspective, mathematical modeling has been used to understand mechanisms promoting and determining cancer evolution^18–22^, but how these mechanisms shape the clonal architecture of a tumor remains largely unexplored.

Here, we integrated mathematical simulations of cancer evolution with algorithmic approaches to infer the clonal architecture of human tumors from their molecular profiles. Precisely, we collected the cancer genomes of 2,673 human tumors from 15 tumor types profiled by The Cancer Genome Atlas (TCGA) (Supplementary Table 1) and infer the clonal composition of each sample from its somatic mutations and copy number alterations^14,23^. Results from *human* tumors were analyzed in parallel with the generation of a large cohort of *simulated* tumors. To this purpose, we used a previously proposed mathematical model of cancer progression^19^ and analyzed the emergence of diverse clonal architectures as a function of alteration and proliferation rates. Results from both types of analyses were highly concordant, validating each other, but also complementary, overcoming the limitations of each approach. Indeed, numerical simulations allowed the exact determination of clonal architectures during tumor expansion and under controlled parameters. On the other hand, results on human tumors allowed to qualitatively examine intra-tumor heterogeneity in the context of distinct tumor types and genomic lesions.

Using this combination of techniques, we explored fundamental features of intra-tumor heterogeneity, such as how many clones can typically be found in a tumor sample, do they have similar sizes or one accounts for most of the cell population, how did they descend from each other, and, finally, what are the evolutionary and molecular determinants of these features. The characterization of complex clonal architectures is a critical first step towards understanding their organizing principles and predicting their emergence.

## RESULTS

To estimate intra-tumor heterogeneity in 2,673 human tumors profiled by TCGA, we used a combination of two algorithmic approaches. First, we use ABSOLUTE^23^ to integrate mutations and copy number changes in each tumor and determine copy number statuses of mutated and wild-type alleles. Then, we used PhyloWGS^14^ to infer the clonal architecture of a tumor from its set of mutations and copy number alterations. To increase the robustness of our results, we estimated the clonal structure of each TCGA tumor sample based on the set of top scoring predictions made by multiple runs of PhyloWGS, each weighted by its likelihood (see Methods). In parallel, we generated a cohort of ~40,000 simulated tumors using a model of cancer evolution governed by two parameters: the alteration rate μ, which is the probability of one cell to acquire a new mutation during replication, and the fitness *s*, which is associated with the probability of a cell to replicate^19^. For each simulation, we tracked the number of clones as well as their size and lineage, thus to reconstruct the exact architecture of each simulated tumor. Simulated tumors were generated using a wide range of alteration rate and fitness values to assess the impact of alteration and proliferation rates on intra-tumor heterogeneity.

### Observed and hidden intra-tumor heterogeneity

At first, we explored how the number of clones varied among human and simulated tumors. Within the TCGA dataset, inferred intra-tumor heterogeneity was highly diverse among tumor types (Fig. 1a and Supplementary Table 1). The rank of tumor types based on their mean number of clones bore a striking resemblance with a previously reported rank based on mutation load^24^, indicating that most mutagenic tumors are on average also the most clonally diverse. Nonetheless, the number of inferred clones in each tumor varied considerably and its correlation with the total number of mutations per sample was overall weak (Pearson’s coefficient = 0.09, Supplementary Fig. 1a). Hence, the number of mutations alone is not sufficient to give an estimate of intra-tumor heterogeneity. Conversely, in our set of simulated tumors, the numbers of clones increased linearly with the alteration rate (Fig 1b and Supplementary Fig. 1b) and could reach values orders of magnitude higher than those observed in the human dataset. In part, this discrepancy could be due to the fact that in each simulated tumor we can exactly count all clones, independently of their size, whereas molecular profiling of human samples captures only a portion of the tumor and only the fraction of alterations present in a sufficient number of cells. Indeed, only mutations with a variant allele frequency (VAF) greater than 1% were retained by TCGA (Supplementary Fig. 1c). To apply a similar filtering criteria to the simulated tumors, we only retained clones with a size (number of cells) corresponding to at least 1% of the total cell population. After applying this filter, the range of the number of clones in simulated tumors became comparable to those of the human cohort (Fig. 1c) indicating that simulated tumors are composed by few large clones (up to ~10 clones) and a wide array of undetectable clones. While multiple factors could limit the growth of newly generated clones^25^, this in silico observation suggests that, in human tumors, filtering mutations with low VAF may result in underestimating intra-tumor heterogeneity

**Figure 1:**
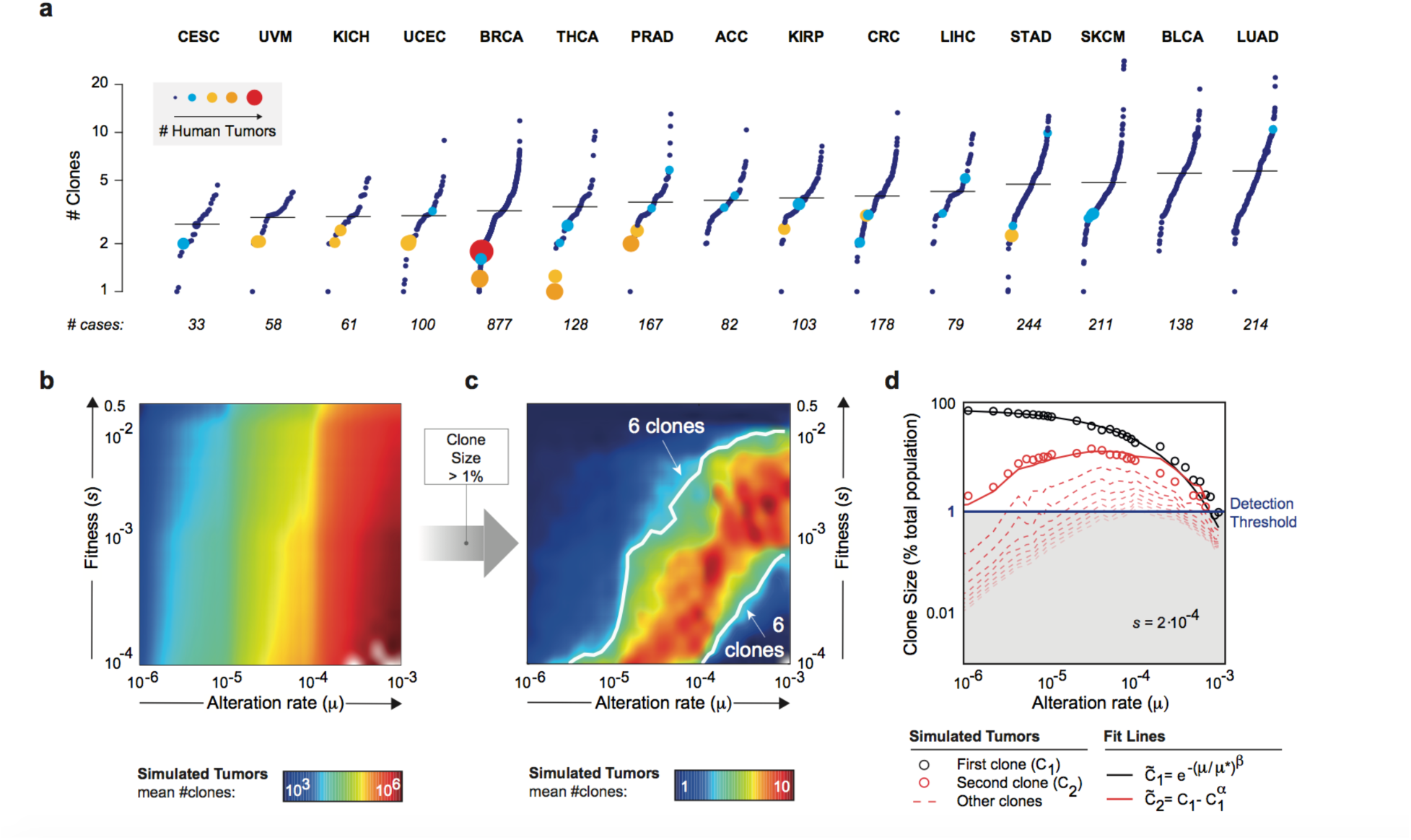
Number of clones in human and simulated tumors. **a)** Number of clones in human tumors within each tumor type. Tumor types are ranked by mean number of clones and tumors within each tumor type are ranked by their respective number of clones. The number of clones in each human tumor is the weighted mean of the number of clones obtained in the top scoring PhyloWGS phylogenies for that sample. Samples with the same mean number of clones are grouped (dots are in size proportional to the number of samples and color coded with increasingly warm colors associated to higher number of clones). **b)** Mean number of clones obtained by simulated tumors as a function of their alteration rate μ (X-axis) and fitness *s* (Y-axis). For each pair of coordinates, μ and s, the mean number of clones observed in simulations corresponding to those coordinates is color coded (cold colors for low numbers, warm colors for high numbers). **c)** Mean number of clones with a size (number of cells) greater than 1% of the total cell population obtained by simulated tumors as a function of their alteration rate μ (X-axis) and fitness *s* (Y-axis). For each pair of coordinates, μ and *s*, the mean number of clones observed in simulations corresponding to those coordinates is color coded (cold colors for low number, warm colors for high numbers). Simulations with 6 clones (white lines) can be found for different ranges of parameters. **d)** The size, as percentage of the total population, of the first clone (black dots) across simulations at different alteration rates and fixed fitness (*s* = 0.0002) decreases with increasing alteration rates μ (X-axis) and is fitted by a stretched exponential (black line). The size of the second clone (red dots) initially grows with increasing alteration rates, but eventually decreases. This trend can be fitted as a function of the size of the first clone and the model parameters (red line - see Methods). Subsequent clones follow the same trend (red dotted lines). Clones with a size greater than 1% (blue line) are detectable. **Acronyms:** CESC (cervical squamous cell carcinoma and endocervical adenocarcinoma), UCEC (uterine corpus endometrial carcinoma), UVM (uveal melanoma), THCA (thyroid carcinoma), KICH (kidney chromophobe carcinoma), BRCA (breast invasive carcinoma), PRAD (prostate adenocarcinoma), KIRP (kidney renal papillary cell carcinoma), ACC (adrenocortical carcinoma), CRC (colorectal carcinoma), LIHC (liver hepatocellular carcinoma), STAD (stomach adenocarcinoma), SKCM (skin cutaneous melanoma), BLCA (bladder urothelial carcinoma), LUAD (lung adenocarcinoma).

The number of detectable clones (i.e. size > 1%) in simulated tumors increased with the alteration rate and decreased with fitness. However, for high alteration rates and relatively low fitness, we observed an unexpected decrease of the number of clones (Fig. 1c, bottom right corner). Before filtering, this decrease was not observed indicating that tumors simulated within this range of parameters were composed by few detectable clones and a large fraction of clones with size below the threshold of 1%. As a consequence, highly heterogeneous simulated tumors now exhibited the same number of observable clones as less heterogeneous ones, “hiding” their true complexity. For example, simulated tumors with 6 clones could now be found for highly diverse alteration rates and fitness and corresponding to tumors that had very different numbers of clones before filtering (Fig 1c, white lines). This result challenges the interpretation of intra-tumor heterogeneity in human tumors, where alteration rates and fitness are unknown features and the amount of undetected clones cannot be estimated.

Interestingly, we observed that the distributions of clone sizes in simulated tumors with high *hidden heterogeneity* were different than those in tumors with low hidden heterogeneity. Indeed, by analyzing the size of the clones in each simulated tumor, we found that the size of the biggest clone decreased with the alteration rate (Fig. 1d). Vice versa, the size of the other clones increased with the alteration rate while μ << s (corresponding to an increase of the number of detectable clones), but as μ approaches *s*, the curves reached a peak and then collapsed below the detection threshold (resulting in a decrease of the number of detectable clones) (Fig. 1d and Supplementary Fig. 2). Also the differences among the sizes of distinct clones in each simulated tumor varied with the alteration rate. At low alteration rates, the first clone is significantly bigger than the second one (Fig. 1d – left side), whereas at high alteration rates their sizes become comparable (Fig. 1d – right side). Importantly, unlike the alteration rate and fitness, clone sizes can be estimated in human tumors. Hence this observation could be used to discriminate between low and high hidden heterogeneity in the human cohort and help estimating the reliability of the inferred number of clones.

### Estimating the hidden heterogeneity in human tumors

To quantify and compare the clone sizes of different simulated and human tumors, we recurred to the concept of population frequency. The population frequency (PF) of a clone *i* corresponds to the fraction of cells in a tumor that exhibit the set of alterations in *i*, but not necessarily only those^14^, i.e. the cells in *i* and in all the clones descending from *i*. Formally, the PF of clone *i* in tumor *T* is defined as:

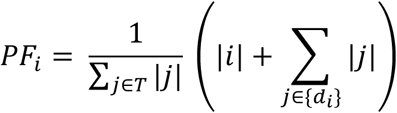

where *|i|* is the size of the clone *i* and *d*_*i*_ is the set of clones descending from *i*. The distribution of sorted PF values (high-to-low) can recapitulate the differences among clone sizes that we observed at varying alteration rates (Fig. 1d). Intuitively, a sharply decreasing distribution indicates that the first clone had time to grow before the emergence of new clones, hence the first clone is considerably bigger than the others (Fig. 2a – green line). Vice versa, a slowly decreasing distribution indicates that new clones rapidly emerged giving rise to detectable clones of similar size (Fig. 2a – red line).

**Figure 2:**
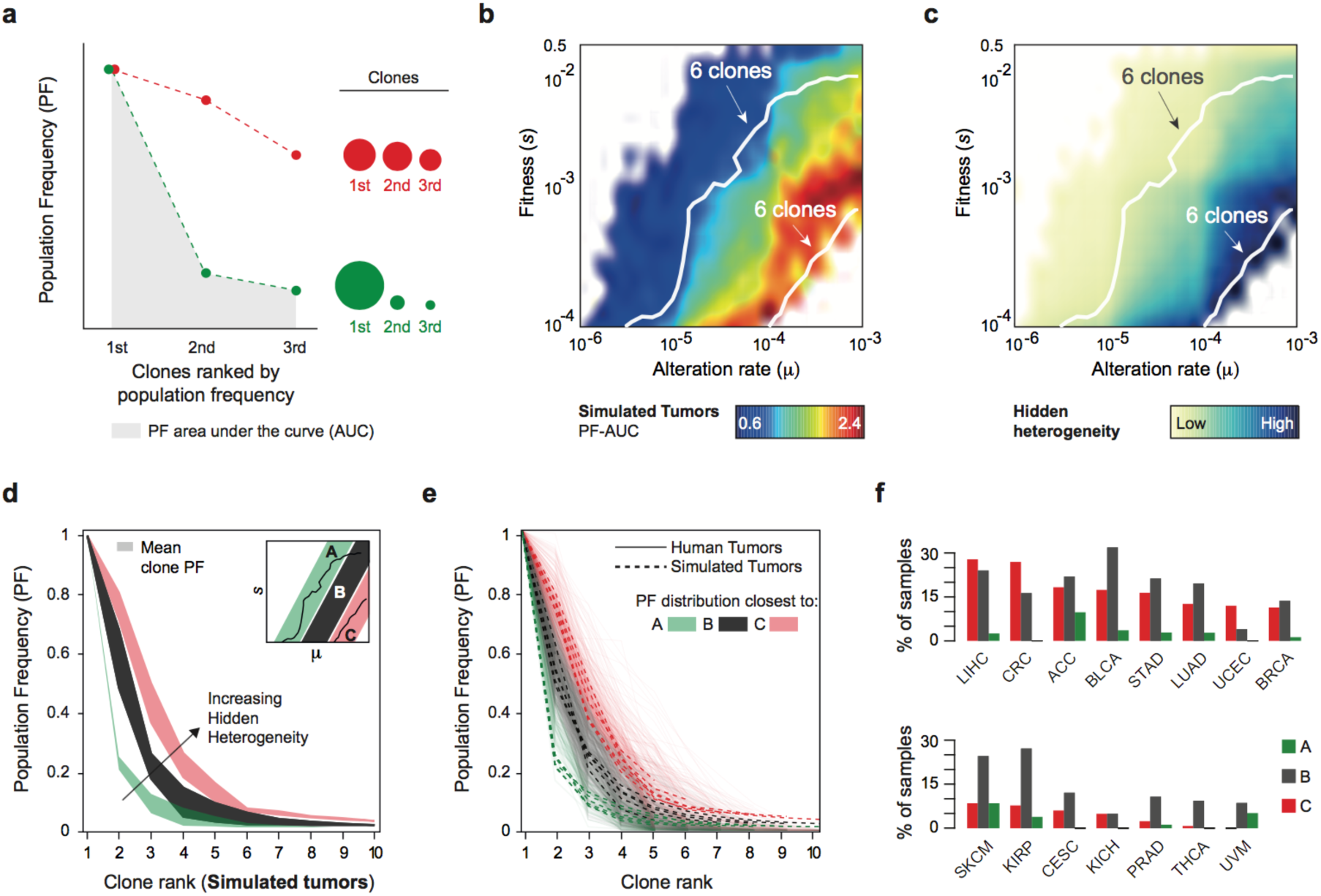
Clone population frequencies predict the extent of hidden heterogeneity. **a)** Schematic of the curves of ranked clone population frequencies. A sharply decreasing curve (green dotted line) corresponds to a large initial clone followed by small clones (green dots with size indicative of the clone size). A slowly decreasing curve (red dotted line) corresponds to clones of a similar size (red dots with size indicative of the clone size). Each curve can be scored by its area under the curve (AUC – e.g. the grey area below the green line). **b)** Clone population frequency AUC (PF-AUC) values of simulated tumors with a number of clones between 5 and 10 as a function of the alteration rate μ (X-axis) and fitness *s* (Y-axis). PF-AUC values are color coded (cold colors for low values, warm colors for high values). Simulation with 6 clones (white lines) have different PF-AUC values based on their range of parameters. **c)** Hidden heterogeneity values (1- fraction of cells in detectable clones) of simulated tumors with a number of clones between 5 and 10 as a function of the alteration rate μ (X-axis) and fitness *s* (Y-axis). Values are color coded (light colors for low values, dark colors for high values). Simulation with 6 clones (white lines) have different extent of hidden heterogeneity. **d)** Ranked clone population frequencies curves for simulated tumors with a number of clones between 5 and 10. PF curves were separately derived for simulated tumors in three ranges of parameters (*μ* and *s*) as shown in the inset on the top right corner (group A in green, group B in black, group C in red). For each group, the corresponding curves were aggregated and the range of values are displayed. **e)** Ranked clone population frequencies curves for human (continuous lines in the background) and simulated (dotted lines on top) tumors with a number of clones between 5 and 10. PF curves of human tumors were assigned to group A, B, or C (lines are colored with the color of the corresponding group) based on the closest curve of simulated tumors with the same number of clones. **f)** Within each tumor type, human tumors were assigned to group A (green bars), B (black), or C (red) based on their PF curves. The percentage of cases assigned to each group in each tumor type is here reported. Only human tumors with a number of clones between 5 and 10 were considered.

In our simulated cohort, we estimated the sorted clone PF distribution in tumors with 5 to 10 clones and scored each simulation by the area under the curve (AUC) of the estimated distribution (Fig 2a – gray area): sharply decreasing distributions will have small PF-AUC values, whereas slowly decreasing distributions will obtain high PF-AUC values. PF-AUC scores increased with alteration rates and decreased with fitness (Fig. 2b). Strikingly, PF-AUC values were highly correlated with the extent of hidden heterogeneity (see Fig. 2b and 2c) and could discriminate among simulated tumors with the same number of detectable clones, but different extent of hidden heterogeneity. Simulated tumors characterized by low hidden heterogeneity (Fig. 2d – region A) had sharply decreasing PF distributions, consistent with the presence of a dominant large clone, vice versa, simulated tumors with high hidden heterogeneity (Fig. 2d – region C) were characterized by PF distributions decreasing slower and, thus, by multiple clones of similar size. A third group of simulated tumors had a higher mean number of clones (see Fig 1c) and intermediate level of hidden heterogeneity (Fig. 2d – region B).

Importantly, unlike alteration rate and fitness, clone population frequencies can be inferred in human tumors and used to estimate the extent of hidden heterogeneity. PF distributions in human tumors were highly consistent with those observed in simulated tumors (Fig 2e and Supplementary Fig. 3). To explore the extent of hidden heterogeneity in distinct tumor types, we assigned each human sample to one of the 3 groups (A, B, and C) by matching its PF distribution to the closest mean distribution obtained in each group by simulated tumors with the same number of clones. Similarly to simulated tumors, human tumor types with the highest mean number of clones (e.g. melanoma, lung, stomach, and bladder cancer) were mostly categorized in group B (Fig. 2f). Instead, colorectal and liver carcinoma samples exhibited the highest percentage of samples categorized in group C, where the estimated hidden heterogeneity is greatest (Fig. 2f). These results suggest that the clonal heterogeneity within these tumor types, or at least within some of these cases might be highly underestimated.

Overall, these results indicate that clone population frequencies and PF-AUC scores can provide a simple way to discriminate among human tumors with similar observed intra-tumor heterogeneity those with greater hidden heterogeneity.

### From clone sets to clonal architectures

To capture the evolutionary history of a tumor, we need to understand how its distinct clones are organized into lineages, i.e. which descend from another. Observed and inferred tumor phylogenies allow to recapitulate this organization. *Linear* phylogenies are the result of the sequential generation of clones along the same lineage, i.e. the last clone is the product and summary of all its predecessors. Vice versa, in *divergent* phylogenies multiple clones spur from the same common ancestor, generating independent lineages that can evolve in clonal populations with little similarity from one another. Tumor phylogenies are typically combinations of linear and divergent evolution and they can be represented as *trees* where clones are the nodes of the tree and two clones are connected if one descends from the other^14^. According to this representation, the first emerged clone is the *root* of the tree, while latest emerging clones are its *leaves.* Intuitively, the more divergent a phylogeny, the closer each leaf will be to the root, in contrast perfectly linear phylogenies will have only one leaf at the maximal possible distance from its root. We formalized this intuition and quantify each phylogeny with the following score:

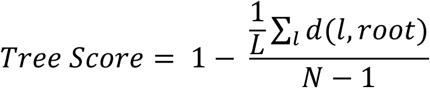

where *L* is the total number of leaves, *N* the total number of clones, and *d(l, root)* is the length of the path connecting a leaf *l* to the *root* of the tree. Based on this definition, all linear phylogenies will obtain a score equal to 0, while the Tree Score will increase with greater branching and number of nodes (Fig. 3a).

**Figure 3:**
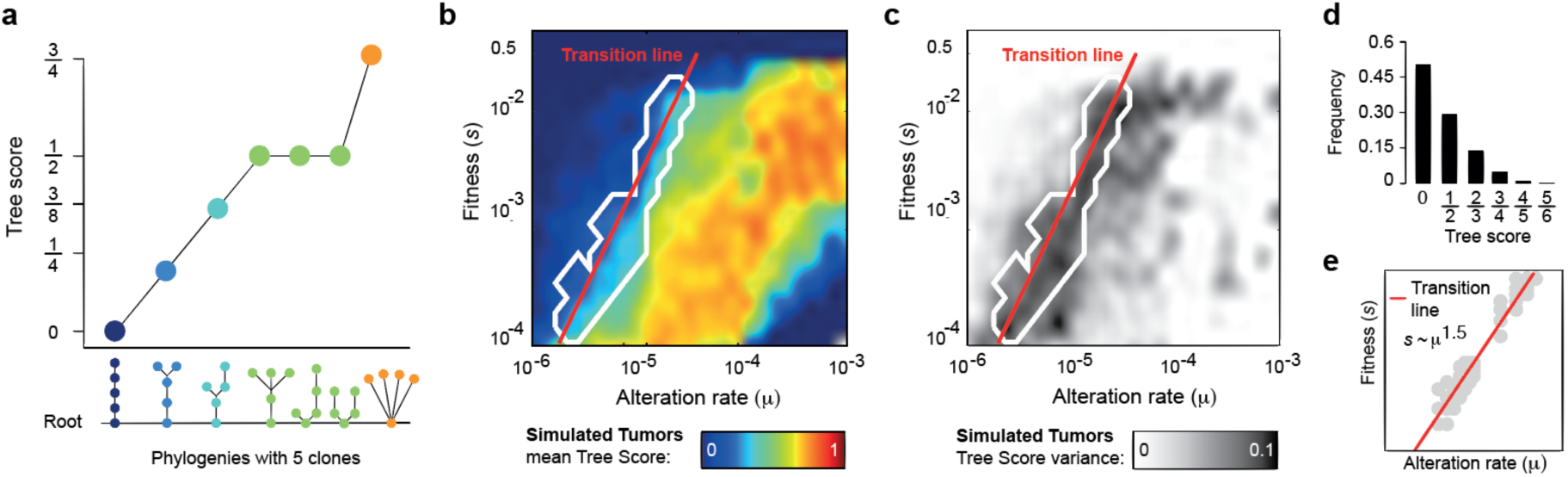
Linear and divergent evolution in simulated tumors. **a)** Example of Tree score values for all possible phylogenies with 5 clones. Tree scores increase with increasing divergence. **b)** Tree scores of simulated tumors as a function of the alteration rate μ (X-axis) and fitness *s* (Y-axis). Tree scores are color coded (cold colors for low scores, warm colors for high scores). The region (white contour) and fitted line (red line) corresponding to the transition between linear to divergent phylogenies are highlighted. **c)** Tree score variance values of simulated tumors as a function of the alteration rate μ (X-axis) and fitness *s* (Y-axis). Tree score variances are color coded (white to black corresponding to low to high variance). The region (white contour) and fitted line (red line) corresponding to the transition between linear to divergent phylogenies are highlighted. **d)** Tree score distribution of tumors within the region of transition. **e)** The transition line (red line) was derived by fitting the points with maximal variance (gray dots).

First, we examined the tree scores obtained by our simulated tumors as a function of both fitness and alteration rate values. The resulting distribution of Tree scores (Fig. 3b) resembled the one observed for the number of clones (Fig. 1c) with more heterogeneous simulated tumors also receiving higher scores. What was peculiar of the Tree score distribution was instead its variance (Fig. 3c). Indeed, we identified two broad areas where simulated tumors were either always characterized by low Tree scores, corresponding to low divergence or linear phylogenies, or always characterized by high Tree scores, corresponding to highly divergent phylogenies. Interestingly, these areas were separated by a narrow region where alteration rate and fitness values allowed the emergence of a wide variety of phylogenies with highly variable Tree scores (Fig. 3d). This result is reminiscent of a phase transition in physical systems, where the variance of observable quantities (called order parameters) diverge at the transition. In our case, the transition is characterized by two phases: linear and divergent evolution. Such a transition indicates that for specific parameters (along the line 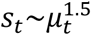 Fig. 3e), linear and divergent phylogenies are equally likely to emerge and cannot be anticipated.

Next, we investigated the association between number of clones and tumor phylogenies. Number of clones and Tree scores were correlated in simulated tumors (Fig. 4a) and as the number of clones increased, these tended to be invariably organized in divergent phylogenies. Surprisingly, multiple clones organized in long linear phylogenies (or phylogenies with minimal divergence) were indeed almost completely absent (Fig. 4a, bottom right corner) indicating that these phylogenies are unexpected. Using the phylogenies inferred by PhyloWGS, we could repeat the same analysis in human tumors. The resulting association between Tree scores and number of clones was remarkably consistent with what the model predicted (Fig. 4b and Supplementary Table 1). Indeed, highly divergent phylogenies were prevalent in human tumors with high number clones, while linear phylogenies with numerous clones were not observed.

**Figure 4:**
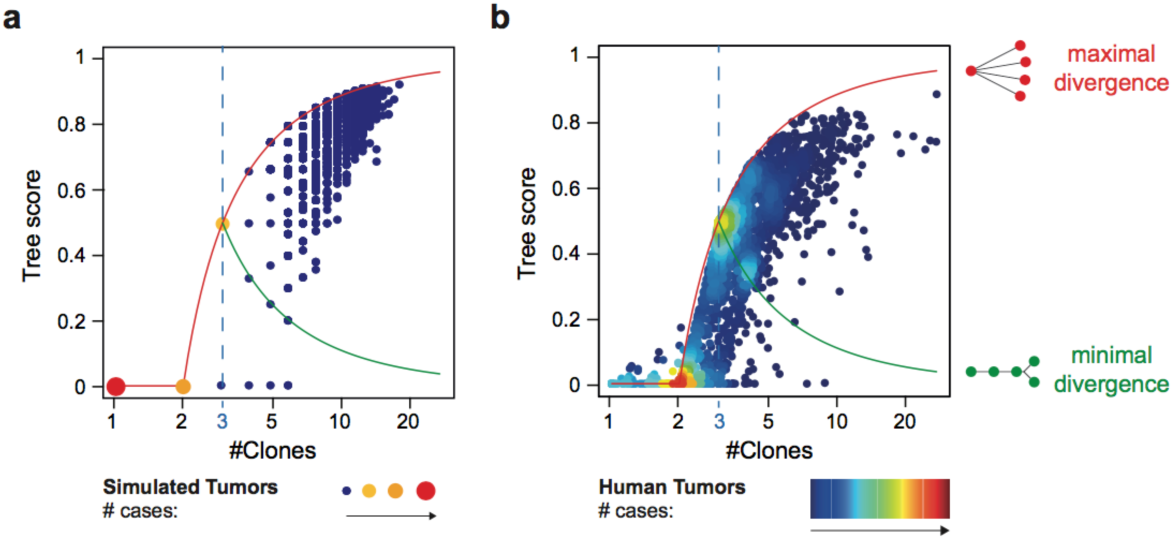
Linear and divergent evolution for low and high number of clones. **a-b)** Tree score as a function of the number of clones observed in simulated (**a**) and human (**b**) tumors. Divergent phylogenies can emerge when at least 3 clones are detected (blue dotted line). The range of Tree scores for phylogenies with more than 3 clones goes from a minimal divergence value (green line) to a maximal divergence value (red line).

As for the number of clones, we observed high variability of Tree scores among human tumors within each tumor type (Supplementary Fig. 4a) and a similarly weak correlation between Tree scores and the overall number of mutations (Pearson’s coefficient = 0.1, Supplementary Fig. 4b). High genomic instability can therefore be observed in both tumors with high and low intra-tumor heterogeneity.

### Clonal and subclonal genomic instability

Previous characterizations of tumor architectures have focused on the dichotomy between clonal and subclonal mutations^16,26,27^: the first are present in all cancer cells and are grouped in the root of the tumor phylogeny (also referred to as the *trunk* of the tree^28^), whereas the second characterize only subsets or individual clones positioned further in the phylogeny. Tumor types in our human dataset exhibited a variable average number of clonal mutations ranging between 40 to 60% of the total number of mutations (Fig 5a and Supplementary Table 1). Highly mutagenic tumors such as lung, bladder, and stomach cancers were on the lower end of this range. Indeed, despite exhibiting a high number of clonal mutations, they were also characterized by numerous subclonal events. A notable exception to this trend was skin melanoma which was characterized by the highest number of clonal mutations, consistent with all of these samples being metastatic and not primary tumors (Fig. 5a).

**Figure 5:**
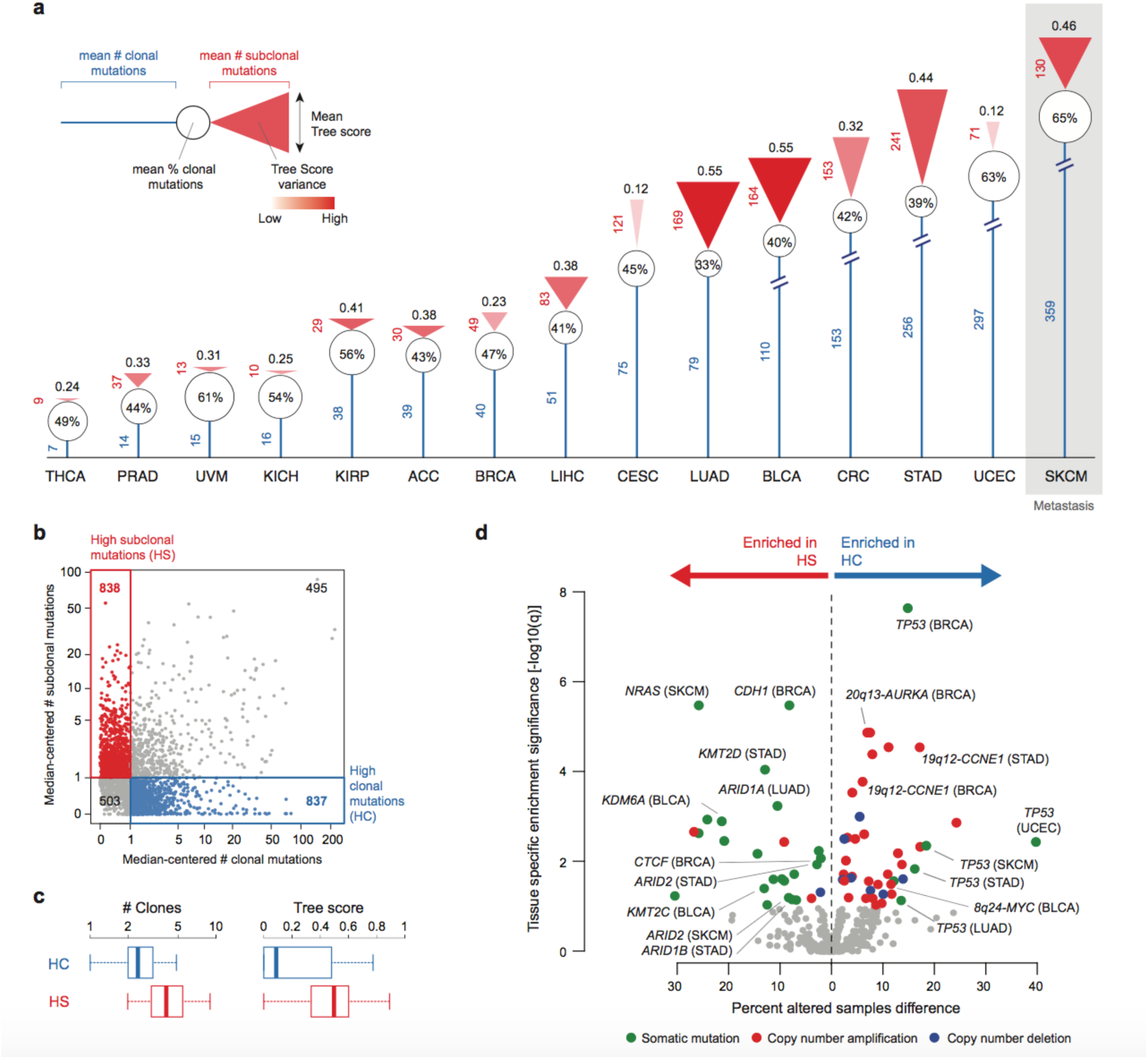
Clonal and subclonal genomic instability. **a)** Mean clonal and subclonal mutations found in each tumor type. For each tumor type we report: mean number of clonal mutations (blue line), mean percentage of clonal mutations (white circle), mean number of subclonal mutations (height of the red triangle), mean Tree score (base of the red triangle), and mean Tree score variance (shade of red within the triangle: intense red corresponds to high variance, transparent red corresponds to low variance). **b)** Median-normalized clonal mutations (X-axis) versus median-normalized subclonal mutations (Y-axis) for all human tumors (dots). The number of clonal and subclonal mutations in each tumor is divided by the clonal and subclonal median value of the corresponding tumor type. Human tumors are grouped into 4 categories based on their numbers of clonal and subclonal mutations being higher or lower than the median of the corresponding tumor type. Samples with clonal mutations higher than the median and subclonal mutations lower than the median are in the High Clonal mutation (HC) group (blue dots). Samples with clonal mutations lower than the median and subclonal mutations higher than the median are in the High Subclonal mutation (HS) group (red dots). **c)** The number of clones and Tree scores of HS human tumors are significantly higher than those of HC human tumors (#clones: Wilcoxon-test p-value = 1.5E-119, Tree score: Wilcoxon-test p-value =1.1E-72). **d)** Volcano plot of cancer-associated alterations tested for enrichment in either HC or HS group. For each alteration we assessed the difference between the percentage of altered samples in the HC group and in the HS group (X-axis) and tested the enrichment in either class by two-tail exact Fisher’s test (negative log10 of the FDR corrected p-values - q-values - are on the Y-axis). Significantly enriched alterations (q-value < 0.05) are color coded based on the alteration type (green: mutation, red: copy number amplifications, blue: copy number deletions), non-significant alterations are in gray. Labels were added to the most enriched in each category and those associated with cell proliferation (for HC enriched events) and with chromatin remodeling (for HS enriched events).

The numbers of clonal and subclonal mutations were nonetheless highly variable among samples, even within the same tumor type (Supplementary Fig. 5a) and showed high fractions of tumors with numerous mutations segregated either in a class characterized by high numbers of clonal mutations (high clonal or HC) or in a class characterized by high numbers of subclonal mutations (high subclonal or HS) (Fig. 5b). Consistently, HS and HC tumors had significantly different number of clones and Tree scores (Fig. 5c), suggesting that genetic features discriminating between HS and HC tumors could also determine of intra-tumor heterogeneity.

To explore whether selected genetic alterations were enriched in either of these classes, we assess the incidence of ~500 cancer-associated somatic mutations and copy number alterations^29^ in HC and HS tumors. Surprisingly, most somatic mutations were enriched in the HS group, whereas copy number changes and *TP53* mutations were most prevalent in the HC group (Supplementary Fig. 5b). To correct for potential associations with specific tumor types, we tested the same panel of alterations within each tumor type separately. *TP53* mutations were significantly enriched in HC tumors from 5 different tumor types (FDR q-value < 0.1). Moreover, in HC tumors we confirmed a higher incidence of copy number alterations across multiple tumor types and in particular of those affecting proliferation-associated genes such as *CCNE1,* Aurora kinase A *(AURKA)* and *MYC* (Fig. 5d). In contrast, HS tumors were found enriched for mutations at multiple chromatin remodelers including SWI/SNF components *ARID1A, ARID1B,* and *ARID2,* the chromatin insulator *CTCF,* and histone modifiers *KMT2D, KMT2C,* and *KDM6A* (Fig. 5d).

High genomic instability can thus be characterized by both numerous clonal and subclonal mutations giving rise to different intra-tumor architectures. Interestingly, specific sets of observed alterations were associated with clonal and subclonal genomic instability. While the role of the altered cellular processes on promoting intra-tumor heterogeneity remain to be elucidated, these alterations could anticipate the clonal diversity of a tumor in a tumor-type independent manner.

## DISCUSSION

Intra-tumor heterogeneity is intrinsically difficult to measure as a limited portion of a tumor is typically accessible for molecular analyses, which provide only a static snapshot of a disease in constant evolution. Computational techniques can help to infer the process that led to tumor formation, extract shared evolutionary patterns through the analysis and comparison of large-scale sample cohorts, and predict the missing pieces of an otherwise incomplete picture. In this study, we integrated algorithmic inference of intra-tumor heterogeneity in human tumors and numerical simulations of cancer evolution. These independent approaches identified remarkably concordant associations among expected number of detectable clones, clone population frequencies, and clonal phylogenies. At the same time, the analyses of human and simulated tumors proved to be complementary approaches to capture both molecular and dynamic features of intra-tumor heterogeneity.

Indeed, numerical simulations allowed to discriminate between the observed intra-tumor heterogeneity, characterized by clones and variants present in a sufficient number of cells to be detected by molecular profiling, and a hidden heterogeneity, that cannot be measured in human tumors. We demonstrate that the extent of hidden heterogeneity is highly variable and it does not correlate with the number of detectable clones. Nonetheless, we found that hidden heterogeneity can be estimated by clone population frequencies, providing a unique opportunity to re-assess inferred clonal diversity in human tumors. It should be noted, that distinct clone population frequencies were observed in simulated tumors with the same number of clones but different values of alteration rate and fitness. This association could prove useful to infer these evolutionary parameters in human tumors.

An association between highly mutagenic tumors and multi-clonal architectures was found in both human and simulated tumors. However, human tumors exhibited great variability within tumor types and the overall correlation between number of alterations and clonal diversity was weak. Indeed, high genomic instability was found both at the clonal and subclonal level and it was associated with different clonal architectures and specific genomic alterations. Alterations associated with increased proliferation were more frequent in tumor with numerous clonal mutations. This feature might indicate the emergence of a highly proliferative clone that end up dominating the cell population. It should be noted, that a similar behavior was also supported by our numerical simulations, where intra-tumor heterogeneity was inversely associated with the fitness parameter. On the other side, we found multiple mutations of chromatin modifiers in tumors with high subclonal instability. In light of the reported association between chromatin remodeling complexes and DNA repair^30,31^, this result suggests that altering these complexes favor the insurgence, viability, and co-existence of multiple clones.

Finally, the approaches here adopted purposefully simplify complex dependences among alterations, clones, and cell types. Indeed, the inference of clonal architecture in human tumors is primarily based on whole exome sequencing of single tumor samples. Multiple samples for each patient, possibly distributed over space and time, would greatly increased the accuracy of such prediction. Similarly, the model we adopted ignores spatial constraints, such as interactions with the microenvironment or mechanical constraints between cells and assumes global and constant parameters in each simulation. Interestingly, experiments with alteration rates varying during a single simulation did not change the trends presented in this work (data not shown). On the other hand, it will be important in the future to explore the effect of cell-cell interaction^32^ and deleterious mutations^20^ on tumor heterogeneity.

Targeted sequencing of cancer-associated variants is empowering clinicians with the ability to tailor therapeutic protocols to the genetic fingerprint of each tumor. These decisions however often rely on a single and potentially incomplete observation. While single-cell sequencing or multiple sampling of the same tumor are still for the most part unfeasible in the clinic, the identification of tumors at “high-risk” of intra-tumor heterogeneity could provide a means to better prioritize patients likely to benefit from additional analysis and profiling. Besides the techniques discussed above, additional investigations could include high-depth sequencing of both DNA and RNA to better identify rare variants^33^, analysis of multiple solid and liquid biopsies with shorter follow-ups^34,35^, and computational analyses to infer intra-tumor heterogeneity. With extended models and a growing availability of data from detailed intra-tumor molecular profiling, it will be possible to provide qualitative and quantitative endpoints to systematically characterize the tumor clonal architecture of each patient.

## METHODS

### Inference of tumor phylogenies: PhyloWGS, numerical procedure and scoring

PhyloWGS is a method to infer evolutionary relationships between clonal subpopulations based on variant allele frequencies of point mutations and taking into account copy number alterations at the mutated loci. PhyloWGS provides in output detailed phylogenies representing the clonal evolution, thus inferring the clonal architecture and not only the clonal composition of each tumor. In particular, PhyloWGS does not provide a unique tree representing the phylogenetic evolution of the tumor, but a number of trees, each scored by its complete-data log likelihood^14^. For each sample, we run 10 inference procedures with different seeds and we kept the 50 trees with the highest complete-data log likelihood for each run for a total of 500 phylogenies for each human tumor. We then sorted all the trees by log-likelihood and kept the top 10% (50 trees) for further analysis. For the reduced list of trees, we assigned a score 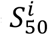 to each tree i according to:

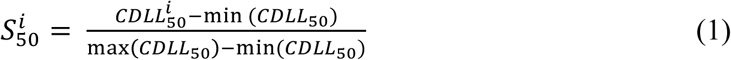

where 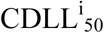 is the complete-data log likelihood of the tree i and min(CDLL_50_) (resp. max(CDLL_50_)) is the minimum (resp. maximum) complete-data log likelihood value within the reduced set of trees. For each sample, we computed the weighted average number of clones and weighted average Tree score as follows:

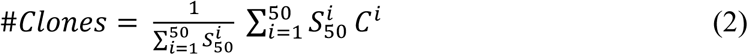

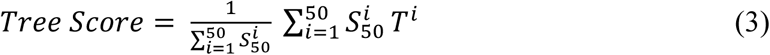

where C^i^ and T^i^ are respectively the number of clones and tree score of the tree i.

### Accuracy of PhyloWGS

PhyloWGS accuracy depends on both the number of mutations and the sequencing read depth. In the original publication, PhyloWGS was applied to synthetic data with known clonal structures to test whether the method was able to recover the true number of clones based on the number of mutations and the read depth. Based on their results, we extract threshold lines for different number of clones in the population separating regions where the reconstruction is accurate and where it is not (Supplementary Fig. 6a-f). For tumors falling above the threshold line, the reconstruction is considered accurate, whereas below the threshold line the number of clones is likely to be overestimated. The vast majority of the TCGA samples we analyzed are in the region of accurate phylogenetic reconstruction. A few cases with high heterogeneity (number of clones > 6) fall slightly below the threshold line indicating a potential overestimation of one clone.

## ABSOLUTE

We used ABSOLUTE^23^ to estimate the copy number status of each point mutation. Originally, ABSOLUTE was designed to infer purity and ploidy of tumor samples, but it also returns information on the copy number status of point mutations when a list of mutations is provided as input. ABSOLUTE reports multiple possible solutions and often manual curation is required to select the best among the top ones (personal communication). For this reason, in this study we relied on TCGA samples with purity and ploidy values previously reported by the authors of the original publication. We independently ran ABSOLUTE on all samples and for each sample i selected the solution that minimizes:

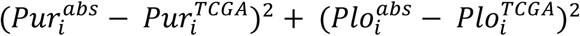

where 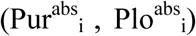 is the purity and the ploidy obtained from our ABSOLUTE runs and 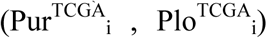 is the purity and the ploidy previously reported for the sample i (Supplementary Fig 6g-h).

### Modeling cancer evolution

To model cancer evolution, we rely on the model proposed by Bozic et. al. ^19^. This model is a discrete time Galton-Watson branching process in which cells can at each time step either replicate (with a probability b) or die (with a probability d). During the replication, one of the two daughter cells can acquire a new alteration with a probability μ. If an alteration occurs, this can be of two types: passenger with a probability μ_p_ and driver with probability μ_d_. The probability μ_d_ is set to 0.025, which corresponds to 500 drivers out of 20000 genes. A driver alterations confers to the cell a selective advantage by reducing its probability to die. The probability to die of a cell i that has accumulated k driver mutations, d^k^_i_ is given by:

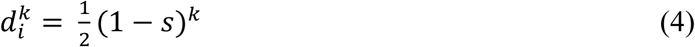

where s is the fitness parameter. According to (1), the replication probability for the cell i with k mutations is b^k^_i_ = 1 − d^k^_i_.μ and s are the input parameters of the model and remain the same during the simulation and for all cells. The probability to die will change during the simulation depending on the number of accumulated driver alterations.

In our analyses, after each replication step, if no alteration has occurred then the two daughter cells will remain in the same clone, otherwise the sibling with the new alteration will create a new clone (Supplementary Fig. 7a). Importantly, a new clone is formed whether the new alteration is a driver or a passenger. To track the full and exact phylogeny corresponding to each simulation is computationally intensive as all cells need to be monitored at each replication step. To improve the efficiency of this computation, we simulate evolution of clones, rather than cells. Specifically, for a clone of size N, the number of replicating cells n_r_ is drawn from a binomial distribution with a success probability b. Then among n_r_, we determine the number of mutating cells n_m_ from a binomial distribution with a probability μ. Lastly, for each altered cell we draw with an equal probability a random number between 0 and 1 and assign a driver (passenger) alteration if the number is lower (higher) than 0.025. Finally, to calculate the mean number of clones and Tree score, only clones with a number of cells greater or equal to 1% of the total population are retained. This is in accordance with the fraction of sequencing reads typically required by cancer exome sequencing studies to retain a somatic mutation (Supplementary Fig. 1). The model of clonal evolution is implemented in Python, using the ETE environment^36^.

### Determination of the size of the clones

Within simulated tumors, we found that the average size of the biggest clone (normalized by the total population size) varies as a stretched exponential with the alteration rate μ (Supplementary Figs. 2a-b):

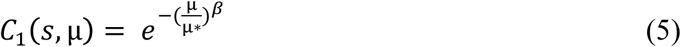

Supplementary Fig. 2 indicates that the characteristic alteration rate μ^□^ increases as a power law with s, μ^□^ □ s^0.821°0.0007^ while the stretching exponent β decreases logarithmically with s as β = −0.37 - 0.028 log(s) (Supplementary Fig. 2c-d). The intervals for the two parameters of the fit are 0.37 ± 0.025 and 0.028 ± 0.003. The fits of μ^□^ and β tend to deviate at large fitness for s > 0.01 from the numerical results, however this regime of very high fitness is not the most relevant for our study since it results in monoclonal tumor evolution.

Since our simulations are bounded by total number of cells (each simulation is stopped once the tumor has reached a size of 500 million cells), the maximal size that new clones can reach is bounded by the size of the largest clone. We determined an expression of the size of subsequent clones that depends on the size of the biggest. Supplementary Fig. 2e shows that we can define the size of clone C_i_ as:

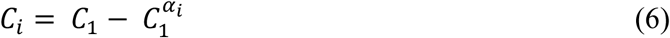

with 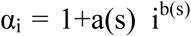 (Supplementary Fig. 2f). Finally, we find a(s) = −0.53-0.022 log(s), with 0.53±0.01 and 0.022±0.0002, (Supplementary Fig. 2g) and b(s) = −1.25-80s, with 1.25±0.007 and 80±4.5, (Supplementary Fig. 2h). Further investigations are required to understand the values of the different constants obtained from the numerical results, but interestingly we observed that several of them are close to the ratio between the probabilities of driver and passenger alterations 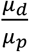□0.025.

Given these expressions for clone sizes we can analytically estimate the number of clones from the values of μ and s. Given μ and s sampled within the space of parameters used in our simulations, we calculated the size of the biggest and subsequent clones using equations (5) and (6). The parameters in the equations are chosen from normal distributions with mean and variance corresponding to the values obtained from the fits. Only clones that satisfied the constraint C_i_ > 0.01 were counted. We repeated this procedure as many times as the total number of simulations and estimate the mean number of clones for each couple of values μ, s. The resulting landscape of analytically predicted number of clones are consistent with the numerically derived (Supplementary Fig. 2i).

### Average ranked population frequency (*PF*) in human samples

To estimate the average ranked population frequency PF, we first sorted the clones inferred by PhyloWGS by their population frequency. For a linear evolution the ranks of the sorted clones are the same as the ranks of the clones from the root to the leaf. However, for phylogenies with multiple branches (or various phylogenies inferred by PhyloWGS with different topologies and/or number of clones) the rank of the clones from the root to the leaves is ambiguous. For this reason, we sorted the clones by their population frequency independently of the underlying topology of the phylogeny (Supplementary Fig. 7b).

After inferring the reduced set of best trees for each human sample and sorting the clones of each tree by their population frequencies, we calculated the average ranked population frequency as:

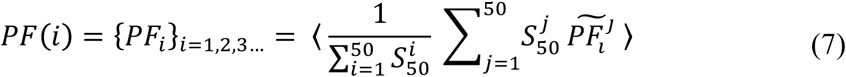

where *i* is the rank of the clone, 〈 〉 denotes the ensemble average over all samples, and 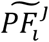 is the population frequency of the clone *i* of the *jth* inferred phylogeny (^~^ indicates that the clones have been sorted by their population frequency in the phylogeny *j*). We should note that to compare the ranked population frequency in simulations and in human samples at a given number of clones (Supplementary Fig. 3), we considered for the human samples the maximum number of clones inferred by PhyloWGS and not the average number of clones. Furthermore, human samples with a maximum PF lower than 0.75 were removed from this analysis, as these tumors are likely characterized low purity (for 100% tumor content, the max PF is equal to 1).

### Alteration enrichment analysis in HS and HC tumors

Tumors with high numbers of clonal (HC) or high numbers of subclonal (HS) mutations were defined within each tumor type as the samples with either a number of clonal mutations higher than the median and subclonal mutations lower than the median (HC), or a number of clonal mutations lower than the median and subclonal mutations higher than the median (HS).

Alteration enrichment analyses were performed both aggregating all HC and HS samples in two pan-cancer groups (Supplementary Fig. 5b) and within each tumor type separately (Fig. 5d). For each analysis, we tested a set of previously characterized 505 cancer-associated genetic alterations^29^. Each alteration was tested if occurring in at least 5 samples of the analyzed dataset. The enrichment of each alteration in either the HS or HC group was statistically tested by a two-tail exact Fisher’s test and all p-values were corrected for False Discovery Rate (FDR) using the Benjamini-Hochberg method.

## REFERENCES

1. McGranahan, N. & Swanton, C. Clonal Heterogeneity and Tumor Evolution: Past, Present, and the Future. Cell 168, 613–628 (2017).

2. Vogelstein, B. et al. Cancer genome landscapes. Science 339, 1546–1558 (2013).

3. Negrini, S., Gorgoulis, V. G. & Halazonetis, T. D. Genomic instability — an evolving hallmark of cancer. Nat. Rev. Mol. Cell Biol. 11, 220–228 (2010).

4. Nowell, P. C. The clonal evolution of tumor cell populations. Science 194, 23–28 (1976).

5. Navin, N. et al. Tumour evolution inferred by single-cell sequencing. Nature 472, 90–94 (2011).

6. Ding, L. et al. Clonal evolution in relapsed acute myeloid leukaemia revealed by whole-genome sequencing. Nature 481, 506–510 (2012).

7. Patel, A. P. et al. Single-cell RNA-seq highlights intratumoral heterogeneity in primary glioblastoma. Science 344, 1396–1401 (2014).

8. Tirosh, I. et al. Dissecting the multicellular ecosystem of metastatic melanoma by single-cell RNA-seq. Science 352, 189–196 (2016).

9. Gerlinger, M. et al. Intratumor heterogeneity and branched evolution revealed by multiregion sequencing. N. Engl. J. Med. 366, 883–892 (2012).

10. Bruin, E. C. de et al. Spatial and temporal diversity in genomic instability processes defines lung cancer evolution. Science 346, 251–256 (2014).

11. Zhang, J. et al. Intratumor heterogeneity in localized lung adenocarcinomas delineated by multiregion sequencing. Science 346, 256–259 (2014).

12. Miller, C. A. et al. SciClone: Inferring Clonal Architecture and Tracking the Spatial and Temporal Patterns of Tumor Evolution. PLOS Comput. Biol. 10, e1003665 (2014).

13. Roth, A. et al. PyClone: statistical inference of clonal population structure in cancer. Nat. Methods 11, 396–398 (2014).

14. Deshwar, A. G. et al. PhyloWGS: Reconstructing subclonal composition and evolution from whole-genome sequencing of tumors. Genome Biol. 16, 35 (2015).

15. Jiang, Y., Qiu, Y., Minn, A. J. & Zhang, N. R. Assessing intratumor heterogeneity and tracking longitudinal and spatial clonal evolutionary history by next-generation sequencing. Proc. Natl. Acad. Sci. 113, E5528–E5537 (2016).

16. McGranahan, N. et al. Clonal status of actionable driver events and the timing of mutational processes in cancer evolution. Sci. Transl. Med. 7, 283ra54–283ra54 (2015).

17. Andor, N. et al. Pan-cancer analysis of the extent and consequences of intratumor heterogeneity. Nat. Med. 22, 105–113 (2016).

18. Michor, F. et al. Dynamics of chronic myeloid leukaemia. Nature 435, 1267–1270 (2005).

19. Bozic, I. et al. Accumulation of driver and passenger mutations during tumor progression. Proc. Natl. Acad. Sci. 107, 18545–18550 (2010).

20. McFarland, C. D., Mirny, L. A. & Korolev, K. S. Tug-of-war between driver and passenger mutations in cancer and other adaptive processes. Proc. Natl. Acad. Sci. 111, 15138–15143 (2014).

21. Beerenwinkel, N., Schwarz, R. F., Gerstung, M. & Markowetz, F. Cancer Evolution: Mathematical Models and Computational Inference. Syst. Biol. 64, e1–e25 (2015).

22. Bozic, I., Gerold, J. M. & Nowak, M. A. Quantifying Clonal and Subclonal Passenger Mutations in Cancer Evolution. PLOS Comput. Biol. 12, e1004731 (2016).

23. Carter, S. L. et al. Absolute quantification of somatic DNA alterations in human cancer. Nat. Biotechnol. 30, 413–421 (2012).

24. Lawrence, M. S. et al. Mutational heterogeneity in cancer and the search for new cancer-associated genes. Nature 499, 214–218 (2013).

25. Burrell, R. A., McGranahan, N., Bartek, J. & Swanton, C. The causes and consequences of genetic heterogeneity in cancer evolution. Nature 501, 338–345 (2013).

26. Jamal-Hanjani, M. et al. Tracking the Evolution of Non–Small-Cell Lung Cancer. N. Engl. J. Med. 376, 2109–2121 (2017).

27. Yates, L. R. et al. Genomic Evolution of Breast Cancer Metastasis and Relapse. Cancer Cell 32, 169–184.e7 (2017).

28. Swanton, C. Intratumour Heterogeneity: Evolution through Space and Time. Cancer Res. 72, 4875–4882 (2012).

29. Mina, M. et al. Conditional Selection of Genomic Alterations Dictates Cancer Evolution and Oncogenic Dependencies. Cancer Cell 32, 155–168.e6 (2017).

30. Watanabe, R. et al. SWI/SNF factors required for cellular resistance to DNA damage include ARID1A and ARID1B and show interdependent protein stability. Cancer Res. 74, 2465–2475 (2014).

31. Sulli, G., Micco, R. D. & Fagagna, F. d'Adda di. Crosstalk between chromatin state and DNA damage response in cellular senescence and cancer. Nat. Rev. Cancer 12, nrc3344 (2012).

32. Marusyk, A. et al. Non-cell autonomous tumor-growth driving supports sub-clonal heterogeneity. Nature 514, 54–58 (2014).

33. Ciriello, G. et al. Comprehensive Molecular Portraits of Invasive Lobular Breast Cancer. Cell 163, 506–519 (2015).

34. Misale, S. et al. Blockade of EGFR and MEK Intercepts Heterogeneous Mechanisms of Acquired Resistance to Anti-EGFR Therapies in Colorectal Cancer. Sci. Transl. Med. 6, 224ra26–224ra26 (2014).

35. Siravegna, G. et al. Clonal evolution and resistance to EGFR blockade in the blood of colorectal cancer patients. Nat. Med. 21, 795–801 (2015).

36. Huerta-Cepas, J., Serra, F. & Bork, P. ETE 3: Reconstruction, Analysis, and Visualization of Phylogenomic Data. Mol. Biol. Evol. 33, 1635–1638 (2016).

